# Exposing the Diversity of Multiple Infection Patterns

**DOI:** 10.1101/058263

**Authors:** Mircea T. Sofonea, Samuel Alizon, Yannis Michalakis

## Abstract

Natural populations often have to cope with genetically distinct parasites that can coexist, or not, within the same hosts. Theoretical models addressing the evolution of virulence have considered two within host infection outcomes, namely superinfection and coinfection. The field somehow became limited by this dichotomy that does not correspond to an empirical reality, as other infection patterns, namely sets of within-host infection outcomes, are possible. We indeed formally prove there are 114 different infection patterns for the sole recoverable chronic infections caused by horizontally-transmitted microparasites. We afterwards highlight eight infection patterns using an explicit modelling of within-host dynamics that captures a large range of ecological interactions, five of which have been neglected so far. To clarify the terminology related to multiple infections, we introduce terms describing these new relevant patterns and illustrate them with existing biological systems. This characterisation of infection patterns opens new perspectives for understanding the epidemiology and the evolution of parasites.

## 1 Introduction

Parasitism has long been considered as a one-to-one interspecific interaction [1]. This vision is gradually changing due to increasing evidence of the multiplicity of genotypes [2] (in its broadest sense, *i.e.* including strains and species) infecting the same species of plant, animal or bacterial host [3, 4, 5, 6]. Though they have indirectly been studied through microbiological experiments for decades [7], multiple infections are nowadays an active research topic in human [8, 9] and veterinary [10] medicine and in phytopathology [11]. This recent shift in focus is stimulated by new technologies that make it easier to detect within-host parasitic diversity (see e.g. [5]).

The current state of the field has two caveats that we wish to address in this paper. The first pertains to terminology: interactions among several parasites and hosts are complicated enough but confusion is added by the proliferation and/or ambiguous use and misuse of terms characterizing them. The second issue pertains to a limited theoretical context that only considers two epidemiological scenarios known as coinfection and superinfection. This has led researchers in the field to try and force any experimental outcome in this binary reference frame. This inevitably results in even more confusion (and frustration). Below, we first expose in more details these two caveats before addressing them. Limiting ourselves to chronic recoverable infections caused by horizontally-transmitted microparasites, we first list all possible epidemiological scenarios using an appropriate formalism. We then show how an explicit model of within-host interactions can help us explore the diversity of these scenarios.

The great diversity of terms related to multiple infections reflects their growing importance in several fields of biological and medical sciences. One can indeed find different names in the literature for the same phenomenon but also the same name for different phenomena, starting with the term ‘multiple infection’ itself. If not used to qualify the clinical diversity of an infection [12], ‘multiple infection’ is often restrained to cases where several genetically distinct parasites simultaneously infect the same host [13] . Such within-host coexistence of distinct parasite genotypes can be referred to using a variety of terms amongst which ‘multiple-strain infection’ [14], ‘multiple genotype infection’ [2], ‘multi-infection’ [15], ‘mixed infection’ [12], ‘mix-infection’ [16], ‘pluri-infection’ [17], ‘polyinfection’ [18], ‘polymicrobial infection’ [19], ‘genetically diverse infection’ [20], but also ‘coinfection’ [21] and ‘superinfection’ [22] (understood as ‘secondary infection’). Conversely, situations where several parasite genotypes present in the population that cannot coexist in the same individual host can be referred to as ‘superinfection’ [23] (here understood as ‘within-host replacement’) or ‘mutual exclusion’ [24].

Confusion in the frame of reference is a common source of ambiguity in biological nomenclature. In botany for instance, ‘hermaphroditic’ can either refer to the flower, to the individual plant or to the species. Although ‘infection’ usually means “the invasion of a host by a microorganism with subsequent establishment and multiplication of the agent” [25], it can denote the persistence of a parasite at any of the following three levels: the cell [26] (in the case of intracellular parasites, mostly used by virologists), the organism [27] (mostly used by clinicians and epidemiologists) and the population [28] (mostly used by ecologists). For the sake of both disambiguation and generality, we here use ‘multiple infection’ in its broadest sense only, that is the circulation of several parasite genotypes in a host population.

Once inoculated (that is introduced) into a host, a parasite genotype either survives or goes extinct. Hence, there are only two different inoculation outcomes per genotype. The outcome may nevertheless depend on the presence of other genotypes inoculated previously or simultaneously and we define a multiple infection pattern as the set of outcomes over all possible inoculation challenges.

Classical epidemiological models such as *S-I-R* (for Susceptible-Infected-Recovered, [29] ) have naturally been extended to multiple infections, usually by allowing for only two different genotypes [30, 28]. The vast majority of these extensions modelled explicitly only the between-host dynamics and made assumptions on within-host processes. Therefore the within host infection pattern was a modelling assumption and in most models it was either coinfection [28, 31], where inoculations are always infectious and thus distinct parasite genotypes coexist in the long term within the host, or superinfection [30, 23, 32], where within host coexistence never occurs but one genotype always outcompetes the others whatever the inoculation order.

The use of the ‘co-’ prefix added to the term ‘infection’ when more than one parasite genotype is involved is now widely used (see references in[33]). A coinfection can originate from a ‘coinoculation’ [34], when the genotypes are introduced simultaneously into a susceptible (uninfected) host and can both develop an infection. It may also originate from the infection of a host already infected by a different parasite genotype that neither excludes nor is excluded by the first one. It seems preferable to see this latter case as a particular coinfection rather than using a different designation such as ‘superinfection’ [35, 22], ‘sequential infection’ [36, 37] or ‘secondary infection’ [38].

The predominance of super- and coinfection as basic assumptions in models has confined theoretical studies of multiple infections into a binary perspective, while few experimental studies have succeeded in settling the nature of the infection pattern for a given biological system [33]. Some models have tried to escape from this dichotomy by nesting an underlying within-host process in the epidemiology [39, 40, 32, 41] but eventually failed to capture other infection patterns than the canonical two. We explain this by the maintenance of two strong assumptions in these models: parasite survival in single infections and virulence-based genotype replacement.

Empirical studies of multiple infections are usually based on coinoculations, the outcomes of which are then compared to single inoculation outcomes. However, the interpretation of these data remains hindered by the lack of a framework identifying patterns specific to multiple infections, patterns that should rely on the combination of all within-host dynamics endpoints and the conditions that lead to them. For instance, Rigaud *et al.* [2] clearly emphasise the direct benefit for empirical parasitology to be gained from theoretical investigations on what they call “multi-parasite hosts” but they restrict the scope of their investigations by considering that the super/coinfection dichotomy captures all the “interactions” that parasites sharing a host may have. The aim of our work is thus to push back the limits of the multiple infection world, while rationally mapping it. To do so, instead of assuming that infection outcomes are a biological constraint of the given host-parasite system (as in the super/coinfection dichotomy), we model them as the aftermath of within-host parasite growth. This allows us to consider novel situations, for instance when two parasite genotypes require each other’s presence to survive, which gives rise to the description of new infection patterns.

Drawing up an exhaustive list of infection patterns can be done by an abstract identification of infection patterns to maps (see section 2). The problem is that the patterns identified using this logical method have no biological support, be it experimental or theoretical. Another approach consists in focusing on within-host dynamics and all the kind of ecological interactions parasites exhibit inside their host: cooperation, exploitation competition, interference competition and apparent competition, both within and between genotypes [42]. In practice, the within-host outcomes are determined by the stationary parasite load each genotype reaches inside the host. Given the number of possible interactions in the model, these are non trivial to predict. They may even depend on the initial conditions, that is here the inoculation context. Indeed, an inoculation is not necessarily followed by an infection as the presence of an already established parasite genotype may prevent the growth of the newly inoculated one.

Here we investigate these within-host dynamics with a general model for two parasite genotypes that allows several types of interactions, especially those through public goods and spite, which are mutually beneficial and mutually detrimental products respectively [43]. After presenting the model and deriving an exhaustive list of the (eight) infection patterns, we provide biological interpretations and illustrations of each of these. Importantly, these infection patterns do not depend on the specific assumptions of the model. It is only the conditions under which they may arise that do. Our typological result thus opens new perspectives to build more general and flexible epidemiological models and study the evolution of infection patterns. It also allows for an unambiguous categorization of empirical multiple infection patterns that should greatly help to clarify the field. The mathematical details of the model and its thorough analysis are provided in the appendix and readers only interested in the biological implications can skip directly to the description of the infection patterns and the Discussion, where their relevance and biological examples are presented without any mathematical notations.

## 2 Infection pattern diversity: a logician’s approach

With two parasite genotypes, there are four qualitatively different host types, depending on the presence/absence of each genotype: the susceptible (uninfected) hosts, the hosts singly infected by genotype 1, the hosts singly infected by genotype 2 and the doubly infected hosts. We hereafter call these types “classes” and label them 0, 1, 2 and 3 respectively. We extend this class formalism to the composition of the inocula as well, which are the set of parasite individuals newly introduced into a host.

Merely asking whether the two genotypes can chronically infect the same host or not greatly under-estimates the complexity of such an apparently simple setting. For instance, let us assume that the genotypes cannot coexist within one host. Both for clinical, epidemiological or evolutionary investigations, it is still worth knowing what the outcome of the inoculation of a class 1 host by a class 2 inoculum is. Indeed, this challenge can result in a class 1 host (genotype 2 is eliminated), or a class 2 host (genotype 1 is eliminated), or a class 3 host (both genotype coexist) or even lead to recovery or death of the host. Knowing this result provides a key argument to initiate (or not) a prophylaxis against genotype 2 from a clinical point of view. It gives necessary information to design the mathematical model that fits the best the epidemiological network. Finally, it accurately describes the selective pressure acting on both parasite genotypes from an evolutionary point of view.

Formally, an infection pattern is the set of outcomes of all possible inoculation challenges. It contains the information about all the host classes and inoculum classes involved (e.g. whether the doubly infected host class exists or not). It also contains additional information because the same host classes can be observed with different sets of outcomes. For instance, doubly infected hosts can result from the inoculation of one or both singly infected host classes but it can also result from the double inoculation of susceptible hosts. Therefore, the diversity of infection patterns for two genotypes cannot be reduced to a raw coinfection/superinfection dichotomy.

How many chronic infection patterns are there for two genotypes only? The answer to this question clearly depends on a myriad of biological details such as parasite transmission mode, host recovery ability and heterogeneity in host sensitivity to parasites. To show that the infection pattern diversity is unexpectedly large, we choose a simple setting where both parasite genotypes are exclusively horizontally transmitted, recovery is always possible one genotype at a time and the host class alone fully determines the outcome of an inoculation challenge. The latter assumption implies that there is a unique inoculation outcome for each couple of host class and inoculum class.With this determinism, infection patterns can be seen as mathematical maps. In appendix A, we show how to formalise and count the infection patterns using appropriate rules, the main one being the necessity for all infected host classes to be reachable from the susceptible host class through a sequence of inoculation or recovery events. After exhaustive map counting, we find that there are exactly 114 distinct dimorphic infection patterns in our simple setting (see Figure S1-1 in the appendix for the whole list).

This heuristic approach reveals and quantifies a previously undocumented diversity of infection patterns. However, even if none of these can be a priori considered as biologically infeasible, most of them appear to be unlikely. In Figure 1, we show three infection patterns that seem uncommon because they contradict usual conceptions about within-host dynamics. In Fig. 1A, both genotypes mutually invade and replace each other in singly infected hosts and cannot coexist in doubly infected hosts. This goes against the idea that if genotype *i* when initially rare in a host is to outcompete genotype *k* (which is initially abundant), then genotype *i* should also outcompete genotype *k* when *i* is initially abundant. A potential negative frequency-dependent process should be expected to drive the dynamics towards a within-host coexistence of both genotypes. Therefore, even if this pattern cannot qualified as impossible, it supposes very particular conditions to exist. In Fig. 1B, genotype *k* cannot infect if it is inoculated after genotype *i* (the transition from state *i* to state 3 is impossible) but it can if it is inoculated before or simultaneously with the other genotype. This goes against the idea that if a stable coexistence of both genotypes is possible, then the within-host interactions should not be so competitive that one genotype cannot grow (even after the other genotype has settled). Finally, in Fig. 1C within-host coexistence is possible but the sequence of acquisition of the genotypes matters. This goes against the idea that if within-host coexistence is possible, then one inoculation of both genotypes should be enough to reach it, while here genotype *i* needs to modify the environment first. These three infection patterns, as many from the 114 we found, depart from usual conceptions of parasite growth but may be biologically explained by parasite-induced change in within-host niches that persists even after the parasite has vanished (a complex process which will not be considered hereafter).

**Figure 1:**
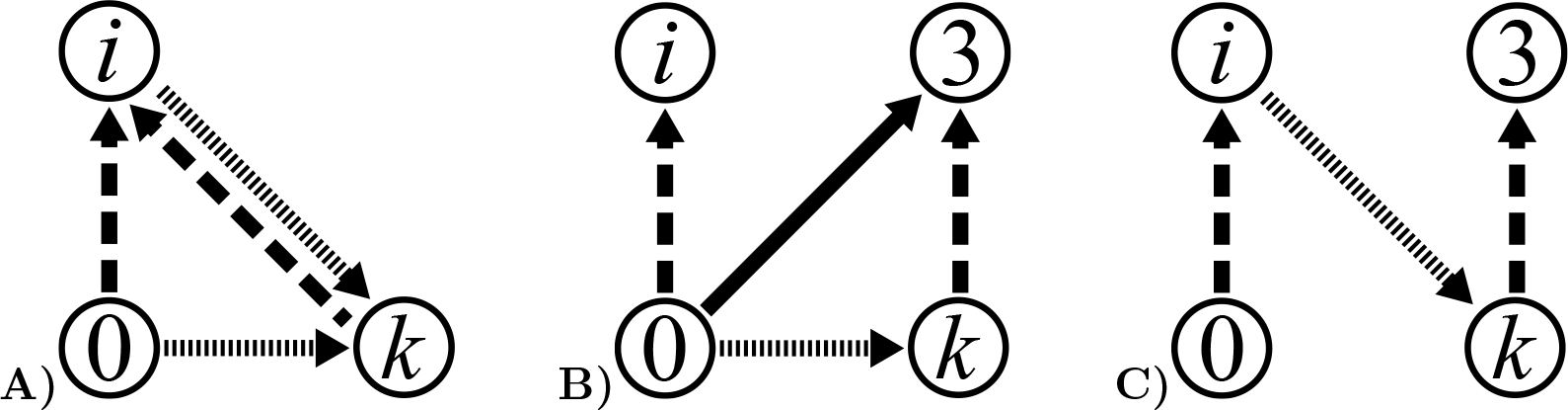
Three potential but uncommon infection patterns found by the exhaustive map counting. A) Mutual invasibility and genotypic replacement of single infections. B) Inoculation order dependent within-host coexistence. C) Infection order dependent within-host coexistence.Circled numbers correspond to host class labels: 0 for susceptible (uninfected) hosts, *i* for hosts singly infected by genotype i, *k* for hosts singly infected by genotype *k* and 3 for hosts infected by both genotypes 1 and 2. Dashed arrows stand for a genotype *i* inoculation, dotted arrows for a genotype *k* inoculation and solid arrows for a 1, 2 double inoculation. For the sake of clarity, looping inoculation and recovery arrows are not shown.

## 3 Within-host modelling

Infection patterns are not all equally biologically relevant, as some of them require the invocation of rare and very specific within-host processes to be observed. In order to determine infection patterns that have an established mechanistic explanation, we develop a general model for the within-host dynamics. Our goal is to refrain from making any *a priori* assumptions on the outcome of an inoculation, that is the long-term persistence of either none, one or both parasite genotypes [44]. This approach allows us to confirm previously described infection patterns but also reveals new patterns, while giving further insight on how parasites interact. To do so we use a modelling framework often used to study microbial communities [45, 46], the competitive Lotka-Volterra model [47, 48].

Here this continuous-time deterministic model is extended by adding public production kinetics. It thus incorporates many types of ecological interactions within and between genotypes (cooperation, exploitative competition, interference competition and apparent competition).

In particular, cooperation and interference competition processes are referred to as public goods and spite production respectively [49]. Inspired by the case of bacteria, we develop a within-host model where public goods correspond to siderophores and spite to bacteriocins. Siderophores are excreted compounds that harvest extracellular Fe^3+^ ions, which are vital for metabolic processes [50]. Provided a receptor compatibility, the iron-siderophore complex can be imported by a bacterium, even if it was not produced by its own species [51]. The empty siderophore is then released back in the environment [52]. As for bacteriocins, they cover a large and diversified family of compounds with a bacteriocidal mode of action [53]. Some of them are fore instance bacteriolytic enzymes that specifically hydrolyse the pentaglycine cross-bridge in peptidoglycan, leading to cell lysis of sensitive strains [54]. Depending on their type, bacteriocins may affect the producing bacteria or not.

We consider two parasite genotypes labelled 1 and 2. For the sake of concision, *i* will hereafter indifferently refer to 1 or 2 and *k* to the other genotype (2 or 1) unless stated otherwise. For a given genotype *i*, *X_i_*, *Y_i_* and *Z_i_* are its parasite load, the concentration of public goods it produces (*e. g.* siderophores [50]) and the concentration of spite molecules it produces (*e. g.* bacteriocins [53]) respectively. These three variables vary with time following a system of ODEs motivated by classical modelling assumptions such as the mass action laws (see [55] for more details): 
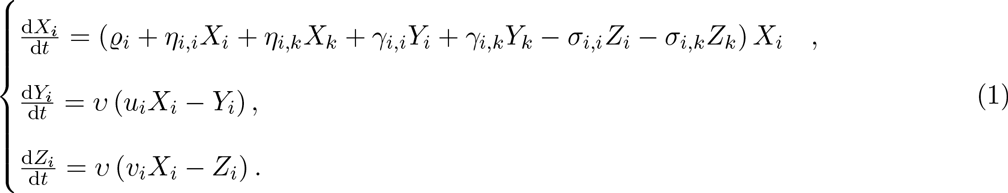

In (1), the lower case Greek letters and *u_i_* and *v_i_* refer to time-independent parameters. Singly indexed parameters denote a given genotype’s trait while doubly indexed parameters denote an interaction effect. In this latter case, the first index refers to the genotype that undergoes the effect, while the second index refers to the genotype from which the effect originates. For self effects, the indices are identical. We denote by *j* and *ℓ* couples of indices that can be identical.

The basic growth rate *ϱ_i_* is the coefficient of the only linear term in the parasite load time derivative. It represents the rate at which the parasite population grows in the absence of all other effects, including self parasite-load dependence. It includes part of the exploitative competition through the parasite efficiency to exploit host resources and reproduce, but also all the costs of constitutive processes such as public goods and spite production.

The parasite-load dependent effects *η*_*j, ℓ*_ are the coefficients of the parasite load quadratic terms in the parasite load time derivative. These include several interactions, namely exploitative competition , apparent competition (through host immune response elicitation and resistance) and direct cooperation. Note that if *η*_*i*,*i*_, then −1/*η*_*j*,*i*_ is the carrying capacity of genotype *i* in the absence of public goods and spite.

The public goods effects *γ*_*j, ℓ*_ are the coefficients of the parasite load - public goods concentration quadratic terms in the parasite load time derivative. They represent the benefits of public goods on parasite growth. The (standardised) public goods production rate *u_i_* weights the secretion of public goods in the host medium. The combination of public goods effects and production rates captures indirect cooperation. Their spite counterparts are denoted by *σ*_*j, ℓ*_ and *v_i_* respectively, and capture interference competition. Finally, *u* is the standard clearing rate of the public productions, which we assume to be independent from their nature or origin. As public productions are often catalytic compounds which are not degraded by their recipient [50, 54], we further assume that the rate at which they are withdrawn from the host medium is essentially due to dilution, self-denaturing, host counteraction or host excretion, that is parasite load independent processes

This model is the adaptation for two genotypes of the within-host part of a general two-level model for multiple infections [55] used to quantify epidemiological feedbacks due to genotype diversity. Here, the focus on the within-host interactions between two parasite genotypes allows us to go deeper in the qualitative analysis of the model and investigate the support for co- and superinfection along with the number and nature of potential other infection patterns. Note that three assumptions made in [55] are relaxed here: self-spiting is possible, parasites can have a negative basic growth rate and a genotype does not always reach the single infection steady state. This increases the diversity of biological situations this model captures. Finally, our within-host model was designed to be as general as possible and, as such, captures several existing models that only include one or two type of interaction between genotypes [56, 13, 57, 58, 44].

The flow diagram in Fig. 2 illustrates the links between variables and parameters according to the set of equations (1).

**Figure 2:**
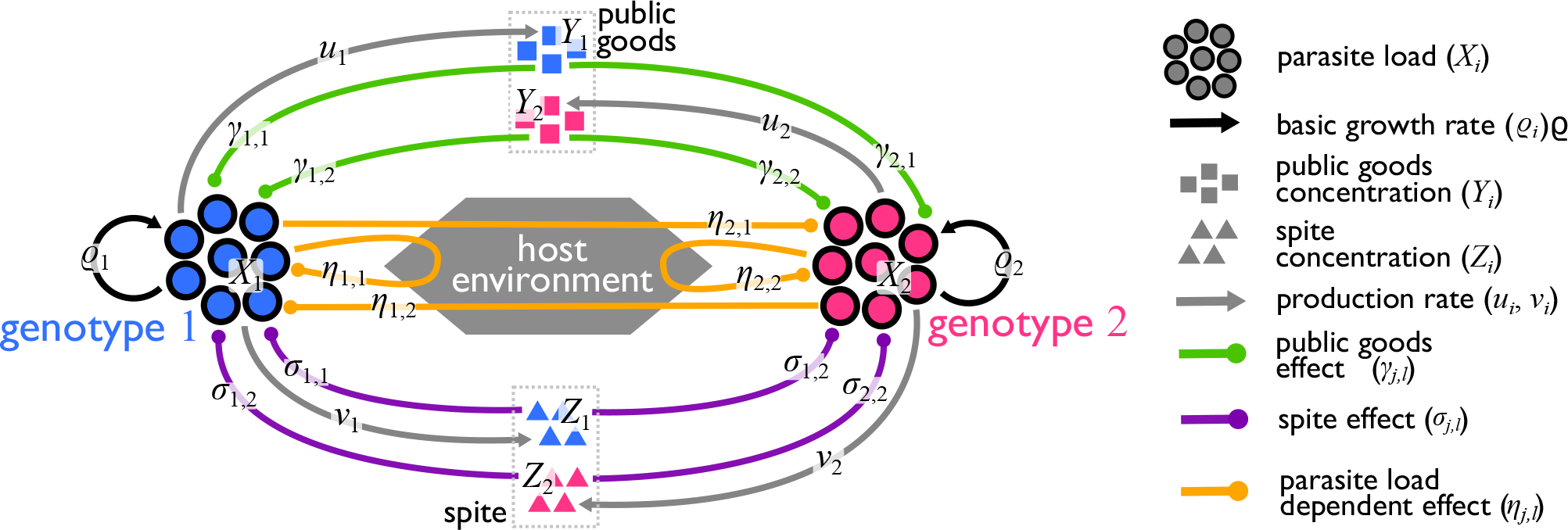
The within-host interaction model. Each genotype produces one kind of public goods and one kind of spite, proportionally to their parasite load. Public productions have beneficial (cooperation - green arrows) and detrimental (interference competition - purple arrows) effects on both genotypes’ growth. The remaining interactions (exploitative and apparent competitions, potentially mediated by the host environment) are captured by the parasite load dependent effects (orange arrows). Note that public production rates are proportional to the same constant *u*, which is the rate at which the public productions are cleared (not shown).

If there is only one parasite genotype in the host, system (1) is characterized only by the three variables related to the inoculated genotype, while it is characterized by six variables when the host is inoculated by both. For this reason, we shall consider two distinct spaces in which (1) can take place (see the appendix B for the vectorial formulation of the model). These spaces are the single inoculation space (SIS) and the double inoculation space (DIS) respectively.

The nonlinearity of system (1) prevents us from finding a general solution. However, the fixed points of the system (i.e. its stationary solutions) can capture its long-term behaviour. For each fixed point, there exists a range of parameters and initial conditions for which the system converges to it. The system has exactly four different fixed points (see appendix C.1 for their derivation): the uninfected fixed point (UFP, where the host is uninfected, none of the parasites can establish an infection), the two singly infected fixed points (SIFP_1_ and SIFP_2_, where the host is infected by only one of the two genotypes), and the doubly infected fixed point (DIFP, where the host is infected by both genotypes). Notice the DIFP is the only fixed point to exclusively lie in the DIS.

Importantly, these theoretical fixed points do not always correspond to biologically relevant situations. A fixed point is biologically relevant if the system can reach it without taking negative values. Mathematically this requires the conjunction of two properties: feasibility and local asymptotic stability (hereafter in the main text termed simply as ‘stability’, but see the appendix for details). Feasibility means that the corresponding steady-state values of the variables are non-negative, while stability refers to the property of the system to converge to this point if close enough at some time-point. If a fixed point is stable, then small perturbations of the system do not affect it on the long run.

Since the variables of a non-inoculated genotype are equal to 0, feasibility in the SIS implies feasilibity in the DIS and vice versa. This is however not true for stability, which can be valid in the SIS and not in the DIS. An interesting biological consequence of this is that the inoculation by genotype *j* in a host already singly infected by genotype *i* can result in the persistence of genotype *j* and in the elimination of genotype *i* from the host.

Feasibility and stability conditions of each fixed point are derived in appendices C.3 and S1-9 respectively. The combination of both conditions (appendix C.5) shows that within-host parameter values govern the outcome of within-host growth in a non-trivial way. The combinatorial complexity that arises when considering each kind of inoculation challenge and asymmetry between genotypes is now to be categorised into infection patterns.

## 4 A biological identification of infection patterns

The previously introduced model provides us a mechanistic insight of the infection patterns it can generate. Interpreting the outcomes of the model at the epidemiological level making Some classical assumptions are needed to link the within-host outcomes with the between-host dynamics but not does not require to specify any model for this greater level, ensuring the generality of our approach. We particularly assume that epidemiological routes of transmission depend only on the long-term behaviour of the within-host dynamics (that is that the within-host parasite growth is fast compared to the transmission/inoculation processes occurring at the between-hosts level). This assumption is common in multilevel models and is justified for chronic infections in which hosts transmit the genotypes that have persisted to the within-host steady state substantially longer than the genotypes that vanished during the transient dynamics [59, 44]. Within-host outcomes can be predicted from the feasibility and the stability of the fixed points of (1). Thus, deciphering all the infection patterns the model can generate is equivalent to characterising all the combinations of fixed points the within-host dynamics can reach.

Before going any further, we need to address a peculiar output of the model when written as (1) , which is explosive infection. Indeed, we show in appendix D that some parameter sets can drive parasite loads to infinity in finite time. One way of dealing with such a biologically unrealistic behaviour of the infection is to assume that there exists a fixed parasite load threshold related to host physiology such that when the total parasite load reaches this threshold all the parasites are killed (see appendix D.4 for the formal modelling). This can either be due to the elicitation of an immune response that kills all parasites or to host death. Similar assumptions have been made in other nested models (see for instance [60]). From an epidemiological standpoint, given our assumption that explosive infections lead to recovery, these have to be interpreted differently depending on the host inoculation status. An explosive infection in singly inoculated hosts should be considered as a non-inoculation because the susceptible host remains susceptible, while an explosive infection caused by a secondary inoculation of singly infected hosts should be considered as a recovery since the initially infecting genotype vanishes.

This rule to handle explosive infections adds to a set of rules that governs the diversity of the sets of feasible and stable fixed points of the three inoculation spaces (SIS_*i*_, SIS_*k*_ and DIS). The feasibility and stability conditions constrain the existence of feasible and stable fixed point sets and also their combination. The set of feasible and stable fixed points of a given inoculation space (the first three columns in Table 1) completely determines the biologically relevant outcome of any host inoculation. The combination of these sets over all inoculation spaces (the lines in Table 1) then determines infection patterns, that is the way both genotypes can circulate through the host population. In appendix E.1, we derive and combine these rules so that all the possible combinations of feasible and stable fixed points sets are explored, leading to the infection pattern list showed in Table 1.

**Table 1:**
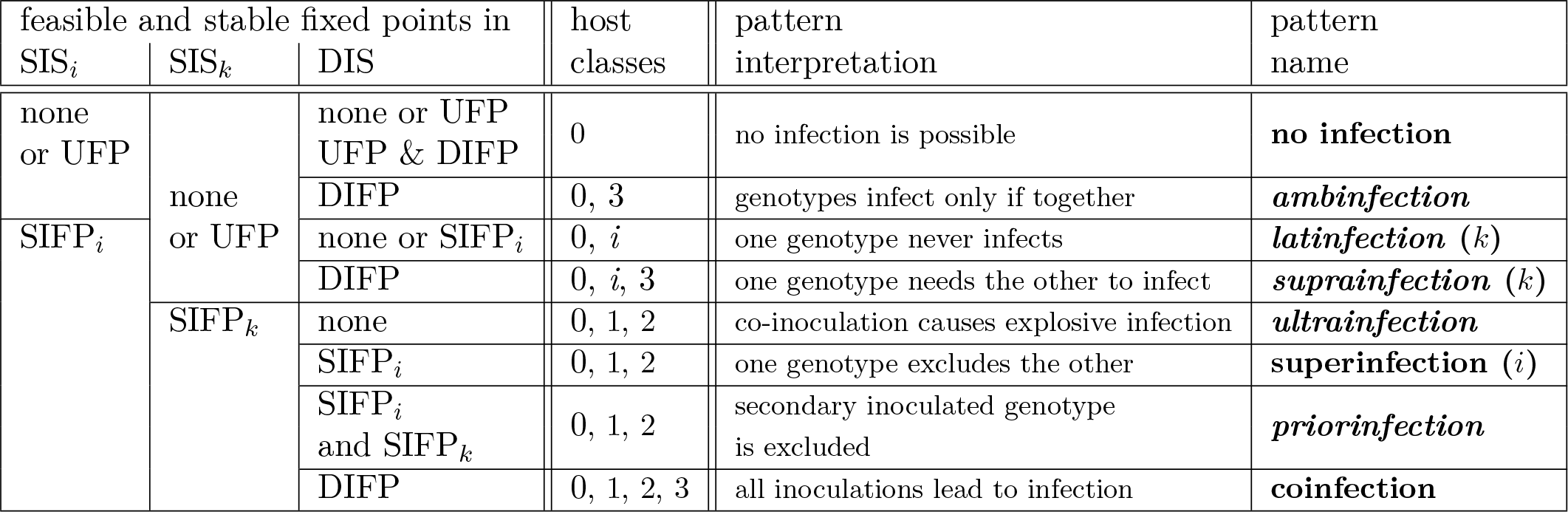
Exhaustive list of infection patterns generated by a multiple infection model, with respect to the sets of feasible and locally asymptotically stable fixed points. Previously unnamed patterns are written in italic letters. For the sake of concision, twin infection patterns (*latinfection i*, *suprainfection i* and *superinfection k*) are not shown.

We find 8 patterns, 5 of which have so far never been studied theoretically. We describe each pattern and suggest an adapted terminology when needed. Links with existing experimental studies are given in the discussion.

**No infection** This pattern arises when no parasite genotype can reach a stable infection state starting from few individuals, neither alone nor in combination. Note that this includes the case where the DIFP is virtually possible but never reached. This pattern is obviously biologically uninteresting and we mention it for the sake of completeness.

***Ambinfection*** This pattern is characterised by the impossibility of single infections: an infection can only originate from a coinoculation of genotypes. We therefore suggest to call it *ambinfection*, from the latin *ambo* meaning “both”.

***Latinfection*** In this pattern, only one parasite genotype is observed, while the other is absent from all within-host outcomes. We call this pattern *latinfection*, from the latin *latere* meaning “to hide”, because the latter genotype is never observed at steady states of infections. Note that we label the two symmetric latinfections according to the apparently absent genotype (“latinfection *i*” thus means that genotype *i* is the “hiding” one).

***Suprainfection*** This pattern is characterised by the impossibility of one genotype to infect without the other. One type of singly infected host is therefore never observed. We call this pattern *suprainfection*, from the latin *supra* meaning “over”, since one genotype can only infect if the other is already present. Note that we label the two symmetric suprainfections according to the double infection restricted genotype (“suprainfection *i*” thus means that genotype *i* is the “extra” one in double infections).

**Superinfection** This well known pattern in ecology and evolution arises when one genotype always out-competes the other in doubly inoculated hosts, whatever the inoculation timings. Note that we label the two symmetric superinfections according to the most competitive genotype (“superinfection i” thus means that genotype *i* always outcompetes genotype *k*). Parasite dynamics within a doubly-inoculated host in a superinfection pattern are illustrated in xsFigure 4A.

***Ultrainfection*** This pattern arises when both genotypes steadily infect singly inoculated hosts but the infection becomes explosive in doubly inoculated hosts leading to an absence of doubly infected hosts. We call this pattern *ultrainfection*, from the latin *ultra* meaning “beyond”, since double inoculation here causes immediate death or recovery. Parasite dynamics within a doubly-inoculated host in an ultrainfection pattern are illustrated in Figure 4B.

***Priorinfection*** This pattern arises when the first genotype infecting a host always excludes the newly inoculated one. Again, doubly infected hosts are never observed. Unlike superinfection, the order of inoculation matters. We call this pattern *priorinfection*, from the latin *prior* meaning “the first of two”, since one genotype can only infect if the other is already present.

**Coinfection** The last pattern is already defined in ecology and evolution. It arises when the two genotypes can infect both alone and together. This is the only case where all four host classes are observed as steady states. Parasite dynamics within a doubly-inoculated host in a coinfection pattern are illustrated in Figure 4C.

**Figure 3:**
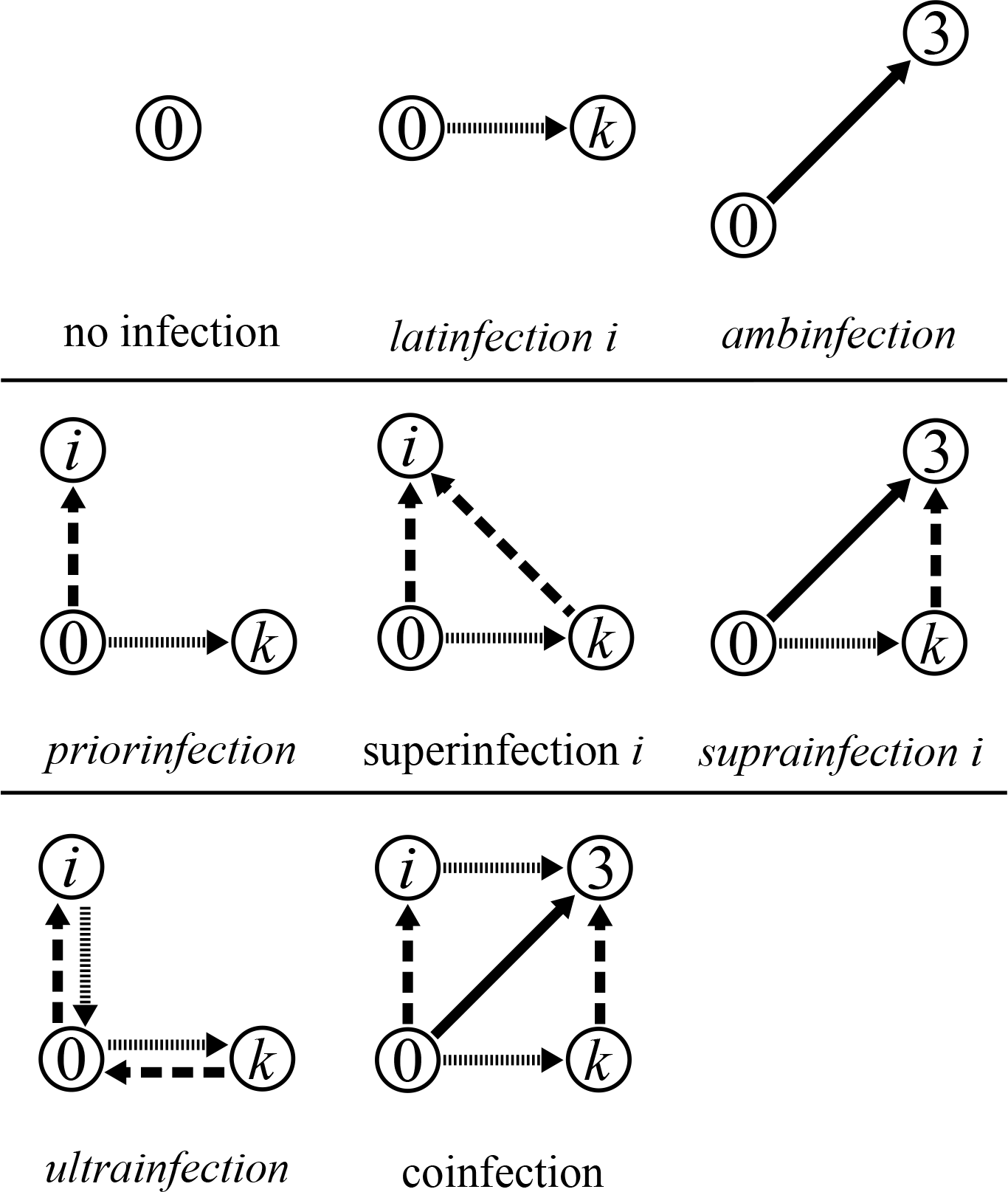
Infection patterns for two parasite genotypes. Note the label-symmetry for the superinfection and *suprainfection* patterns. Circled numbers correspond to host class labels: 0 for susceptible (uninfected) hosts, *i* for hosts singly infected by genotype *i*, *k* for hosts singly infected by genotype *k* and 3 for hosts infected by both genotypes 1 and 2. Dashed arrows stand for a genotype *i* inoculation, dotted arrows for a genotype *k* inoculation and solid arrows for a 1, 2 double inoculation.

**Figure 4:**
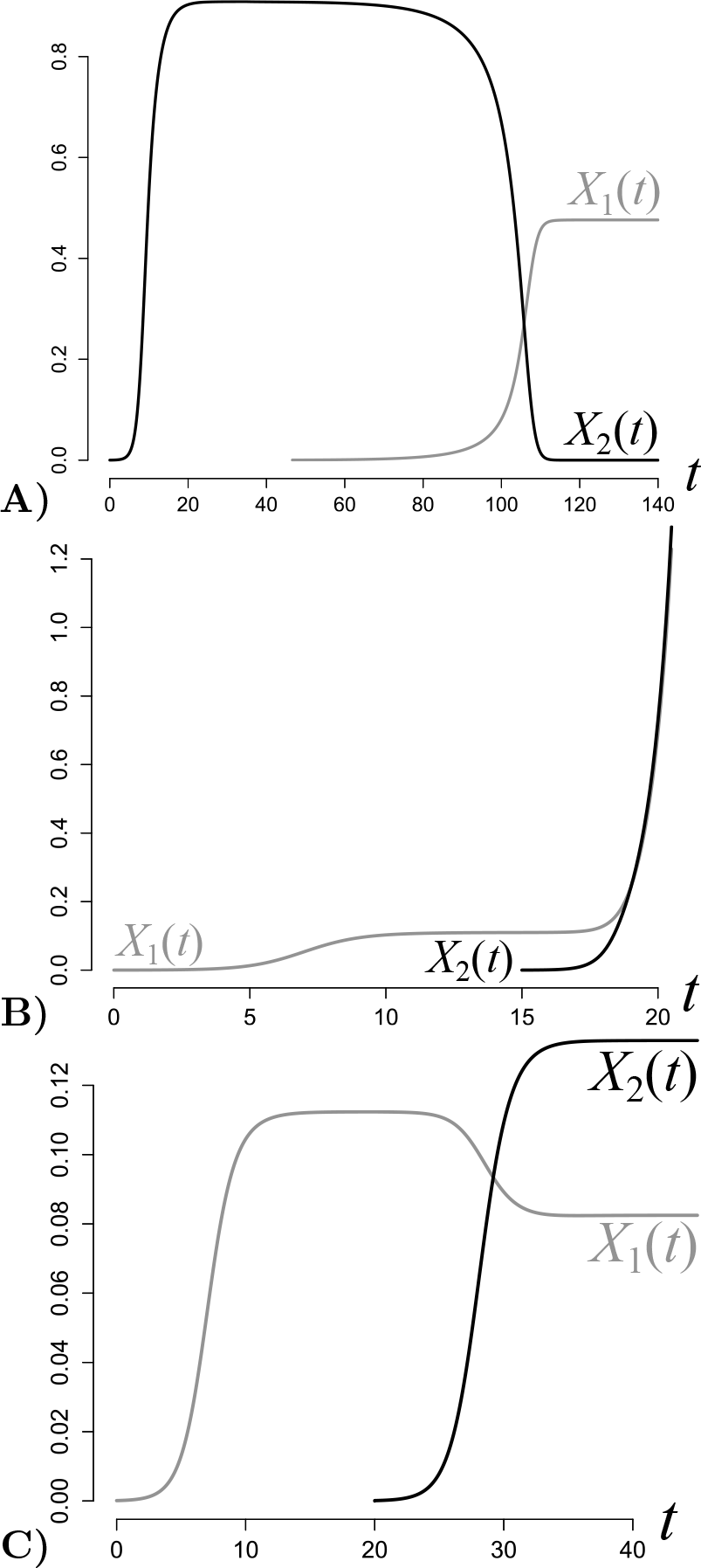
Within-host dynamics of parasite genotype loads in superinfection (A), coinfection (B) and ultrainfection (C). These are the results of numerical integrations of (1) in *R* [61] using the *lsoda* function from the *deSolve* package [62]. **A) Superinfection 1.** At *t* = 0, a host is inoculated with 10^−4^ PLU (parasite load units) of genotype 2. At *t* = 45, the parasite load of genotype 2 (X_2_, in back) has reached its stationary value 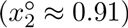 and 10^−4^ PLU of genotype 1 are introduced (*X*_1_ in grey). At *t* = 150, genotype 2 has almost vanished (*X*_2_ has fallen below 10^−6^ PLU) while genotype 1 has reached its stationary value 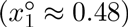 and so even if 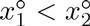. Parameter values are *ƞ*_1,1_ = *ƞ*_2,1_ = −3, *ƞ*_2,2_ = *ƞ*_1,2_ = –2, *γ*_2,1_ = 0.5, *γ*_1,2_ = *σ*_2,1_ = 2, *σ*_1,1_ = *σ*_2,2_ = 0.1 and the others are equal to 1. **B) Ultrainfection.** At *t* = 0, a host is inoculated with 10^−4^ PLU (parasite load units) of genotype 1. At *t* = 15, the parasite load of genotype 1(*X*_1_, in grey) has reached its stationary value 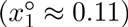 and 10^-−^ PLU of genotype 2 are introduced (*X*_2_ in black). Five time units later, the total parasite load has increased over tenfold. All the parameters are equal to panel A except = *σ*_2,2_ = 0.1, *σ*_1,2_ = 02,*σ*_2,1_ = 0,5, *ƞ*_1,1_ = *ƞ*_2,2_ = −10, *η*_1,2_ = 9, *η*_2,1_ = 10. **C) Coinfection**. At *t* = 0, a host is inoculated with 10^−4^ PLU (parasite load units) of genotype 1. At *t* = 20, the parasite load of genotype 1 (*X*_1_, in grey) has reached its stationary value 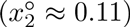 and 10^−4^ PLU of genotype 2 are introduced (*X*_2_ in black). At *t* = 45, both genotype steadily coexist at their stationary values (*x*_1_ ≈ 0.08 and *x*_2_ ≈ 0.13). All the parameters are equal to panel A except *η*_1,1_ = −10, *η*_2,2_ = −8, *η*_1,2_ = −4, *η*_2,1_ = −3, *σ*_1,1_ = *σ*_2,2_ = 0.1.

Finally one can easily remark that any deterministic model assumed with global stability (that is in each space there is at most one fixed point that attracts all trajectories) would not generate other infection patterns than these (and priorinfection would not be generated at all for it requires that two fixed points are local attractors in the same space).

## 5 Discussion

The great number of fields for which multiple infections are a subject of interest, added to time and spatial scale confusions, has lead to a diversified but ambiguous terminology. This terminology essentially relies on the existence (or not) of multiply infected hosts, which is problematic. For instance, the absence of detection of coinfected hosts in a natural population does not imply that parasite genotypes cannot coexist within the same host because it can also be due to between-host limitations such as low contact rates, high recovery rates or high death rates. Another reason why the absence of doubly infected hosts is not sufficient to define superinfection is because an already infecting genotype could exclude the newly inoculated genotype with no absolute competitive hierarchy between genotypes (i.e. a precedence dominance situation). Thus coinfection, superinfection, and what we more generally call “infection patterns”, are within-host level specific terms that cannot be directly inferred from between-host observations. Nevertheless, empirical and theoretical epidemiology of multiple infections rely on knowing the appropriate pattern for determining what must be measured and modelled respectively. Indeed, avoiding a fully open model greatly reduces the complexity of the investigated system in terms of number of expected host classes, compartments, flows and parameters thus allowing for a greater focus on the relevant processes (but see [55] for dealing with this combinatorial complexity).

Modelling epidemiological phenomena, *e.g.* the spread of an infectious disease or host-parasite co-evolution, is essentially about determining the appropriate equations that well describe the dynamics of these phenomena; often these can be seen as flows between compartments. It is obvious that the flows that govern the between-host dynamics are intrinsically related to the infection pattern. Our logician’s approach however reveals that, even in the simple case of two genotypes, the diversity of infection patterns exceeds by far the dichotomy between coinfection and superinfection that most epidemiological models have considered. Designing a different framework for each of the more than a hundred infection patterns, however, would quickly lead to an impractical pile of models, espcecially since many of the potential infection patterns are biologically unlikely to occur. The modelling initiative must thus go beyond the choice of the infection pattern and consider the processes that generate it, that is the underlying within-host dynamics. We therefore advocate that nested modelling, which considers both within and between-host dynamics, is the most appropriate theoretical approach to understand epidemiological phenomena involving parasite diversity, because it provides a unified framework that is able to generate a biologically relevant diversity of infection patterns.

The model presented here captures several within-host interactions between microparasites, notably cooperation and interference competition through explicitly modelled enzymatic productions. It highlights seven non trivial infection patterns, illustrated in Figure 3, that are thus supported by common microbiological mechanisms. Consequently, we have shown that the super/coinfection dichotomy is easily overcome without making any *a priori* assumptions. We show below that the typology exposed in Table S1-10 accurately captures different within-host processes realised in natural biological systems. It is also worth noting that the list of infection patterns is exhaustive for this model: incorporating other processes than the ones considered here (e.g. types of interactions among parasite genotypes) in the model would not alter the number of infection patterns; only the conditions of their existence.

**Coinfection** Two parasite genotypes define a coinfection when they can infect together *and* alone. More precisely, these genotypes are able to cause single infections in distinct hosts and double infections when inoculated within the same host. In the literature, ‘coinfection’ is indeed almost always defined as the “stable coexistence of different parasites in the same hosts” [31]. This leaves the outcome of single genotype inoculation in susceptible hosts unspecified. Yet, all theoretical works assume the effectiveness of single infections for each parasite [31, 28, 63, 40, 41, 64]. Therefore, our definition of coinfection clarifies a previously implicit condition. At the same time, it enables the unambiguous definition of other patterns that share the double infection condition but not the single infection one (*i.e.* suprainfection and ambinfection).

Some experimental articles apply the adjective ‘coinfected’ to hosts that are infected by potentially non self-sufficient parasites, such as satellite viruses (see the discussion part on ambinfection below and [34]). These studies are usually interested in the interactions between parasites and search for synergistic or antagonistic effects that may affect epidemiological traits such as virulence or transmission [65] . However, they do not focus on qualitative properties of the outcome, such as one genotype being rescued by the other. This confusion between models that assume the single infection condition and experimental studies that do not [66] is an additional reason to clarify the definition of ‘coinfection’ and its derived words. As a consequence, when dealing with two genotypes within the same host without information about single infections, we recommend to use the term ‘double infection’.

By analysing the parameter conditions that lead to the coinfection pattern, we find that it occurs when each genotype inhibits its own growth more than that of the other. This is similar to the famous ecological principle of limiting similarity [67]. Nonetheless stationary coexistence within the host does not imply complete neutrality between genotypes. Indeed, as shown in Fig 4 (middle), parasite loads can be impacted positively or negatively by the presence of the other genotype, which can have dramatical consequences on the total parasite load and the overall virulence (but see below and [33]).

In nature, coinfection is a common pattern: *Mycobacterium tuberculosis* and the *Human immunodeficiency virus* (HIV) in humans [68], or the two strains of the *Tomato yellow leaf curl virus*, TYLCV-Mld and TYLCV-IL, in tomato plants [65], are two examples of coinfections, since parasites can infect their host both together and alone. Furthermore, results from metagenomics suggest that coinfection is commonplace [69].

**Superinfection** Superinfection occurs when two parasites can infect alone *and* one always outcompetes the other when together (independently of the order of the inoculations). In the literature, ‘superinfection’ often implies “the exclusion of the less virulent strain” [64, 23, 70]. Note that, as for coinfection, the single infection condition is usually implicit. The fact that it may not be satisfied makes the latinfection pattern possible (see below). Contrary to ours, the definition found in the literature relies on a trait, usually virulence, that characterises each genotype. In its broadest sense, virulence is a shared trait between the host and the parasite and is measured as the host’s fitness loss due to infection [71]. Since there are no doubly infected hosts in a superinfection setting, virulence is necessarily measured in single infections. This is problematic because how could the comparison between two environment-dependent traits, quantified in two distinct environments (the singly infected hosts), rule the outcome of a competition that takes place in a third environment (the doubly inoculated host)? This question still holds if the hosts are clonal, because the parasites themselves shape their environment. The nonlinearity of these processes, including apparent competition, makes the infection outcome of the doubly inoculated host impossible to predict just from virulences quantified (whatever the way) in single infections. Our explicit modelling of the within-host dynamics allows us to overcome this problem and to define superinfection as the exclusion of the same genotype in any context.

Looking at the parameter conditions that lead to superinfection reveals that both genotypes need to inhibit their own growth and that only one of the two genotypes needs to inhibit its growth more than that of its competitor. This asymmetry generates the exclusion of the genotype that somehow competes more with itself than with its competitor. This confirms that the exclusion of a genotype has more to do with their interactions than with their own growth rate, which is related to host resource exploitation and therefore, potentially, to virulence.

To further illustrate the risk of defining superinfection through virulence, let us first make the assumption that, for a given parasite genotype, a greater parasite load decreases host fitness. Let us further assume that for two similar genotypes, the relationship between parasite load and virulence approximately follows the same monotonically increasing function. Figure 4 (left) shows that it is possible for a less virulent genotype to outcompete a more virulent genotype, even when inoculated afterwards. As a conclusion, our model and definition of superinfection capture a greater diversity of asymmetric exclusions freed from the assumption that virulence and competitive ability are positively correlated.

The evidence of competitive exclusion in doubly inoculated hosts has been shown for instance for two strains of the rodent malaria *Plasmodium chabaudi* in mice [72, 73], where it has been reported that secondary inoculated strains decreased the resident strain loads up to six fold and below detectable levels.

**Priorinfection** Priorinfection occurs when both parasite genotypes infect alone *and* when the first genotype inoculated always excludes the other. Unlike superinfection, the infection outcome in a doubly inoculated host depends on the sequence of the inoculations (i.e. the excluded genotype is not always the same).

This pattern is reached if each genotype inhibits its own growth less than it inhibits that of the other. A biological interpretation of this result is that the first genotype in a host modifies the within-host environment, for instance through high spite concentrations of immune system elicitation, so much that any newly inoculated genotype rapidly goes extinct.

The impact of priority on the within-host infection outcome has been noticed for a long time. In 1945, Delbruck showed that *Escherichia coli* (strain “B”) infected with either *a* or *S* bacteriophages cannot be infected by the other phage if inoculated within four minutes after the first inoculation. He called this the “mutual exclusion effect” [24]. The use by plants of RNA interference as an antiviral defence mechanism, spite in our model, notoriously gives rise to priorinfection of plant viruses, sometimes referred to as “cross-protection” [74] or “super-infection exclusion” [37]. Finally, priorinfection echoes vaccination as it prevents secondary infections.

**Ultrainfection** Two parasite genotypes define an ultrainfection when they can infect alone *and* provoke explosive infection when together. Ultrainfection is due to excessive cooperation between the two genotypes. High concentrations of highly beneficial public goods and/or cross facilitation through the alteration of the host’s immune system compensate for the negative density-dependence, which induces explosive infections. In this case, double infections are transient and do not contribute much to the epidemiological dynamics; this may explain why this pattern has been neglected so far. Two honeybee parasites illustrate this pattern: the parasitic mite *Varroa destructor* indeed triggers the replication of the initially cryptic *Deformed wing virus* (DWV) within the bee up to lethal levels, while none of these parasites kills their host alone [75].

More generally, this pattern should be regarded as a limit case of coinfection, where strong cooperative interactions destabilise the within-host dynamics so that parasite loads rise to levels way greater than the ones reached in single infections.

**Latinfection** Latinfection first requires that one genotype succeeds in infecting hosts alone, while the other fails to do so. Second, in doubly inoculated hosts, the former genotype does not help or actively excludes the latter or both grow beyond the total parasite load the host can bear, as the conditions in Table S1-9 suggest (the associated inequality involves trait values of both genotypes). This should not be seen as a classical “single infection” because two kinds of parasite genetic diversities would be missed: transient diversity at the within-host level (both genotypes are present before reaching the single infection fixed point, or the load threshold) and cryptic diversity at the between-host level (if the second genotype appears from deleterious mutations).

Because of lack of both interest (cryptic genotypes have no epidemiological impact) and data (transient within-host diversity is not easy to detect), this pattern is poorly documented. Yet the case of the *Cucumber mosaic virus* (CMV, *Bromoviridae* family) that infects tomato, zucchini squash and muskmelon is a possible illustration of latinfection. Because of deletion and recombination, this RNA virus generates defective RNAs which are not able to infect systemically [76].

Latinfection finally plays a major role in parasite evolution, since it corresponds to stable monomorphism, thus denoting purifying and stabilizing selection.

**Suprainfection** Suprainfection is an asymmetric pattern where one genotype successfully infects the host only in the presence of the other. It captures a diversity of microbial systems and evolutionary scenarios.

This pattern occurs if one of the two genotypes does not cause single infections, either because parasite load explodes or because it vanishes. In the first case, suprainfection occurs only if the second genotype slows down the growth of the first when they come about within the host. The second case is more familiar and compelling because if the first genotype fails to reproduce alone, then suprainfection only occurs if it benefits from the presence of the so-called self-sufficient genotype [77]. This is the cooperative matching piece of latinfection, with the self-sufficient genotype facilitating or being exploited by the other one instead of outcompeting it.

The biology of virus-dependent entities (or subviral agents) is typically based on these two relationships between parasites and thus provides many examples of suprainfection. There are two kinds of virus-dependent entities: virus satellites [78] (which include virophages [79]), which are unrelated to their ‘helper’ virus, and defective viral particles [80], which are replication and recombination byproducts of a parent virus from which they derive. In both cases the helper/parent virus drives at least the replication of the virus-dependent entity, thus allowing it to infect the host as well.

For instance one can find tomato plants doubly infected by the begomovirus *Tomato leaf curl virus* (ToLCV) and its satellite (ToLCV-sat) but the latter is not required for ToLCV infectivity, while it depends on ToLCV for its replication and encapsidation [81]. In 2001, defective dengue-virus type 1 (DENV-1) strain originated from a stop-codon mutation in the surface envelope (E) protein gene. This strain nevertheless achieved long-term transmission in nature thanks to complementation with functional DENV-1 strains within both humans and *Aedes* mosquitoes [66]. Lastly, a well-known example of suprainfection is hepatitis delta virus (HDV), a defective RNA virus the virion coat of which is the one of hepatitis B virus (HBV), thus making HDV unable to infect a human if HBV is not already established or co-inoculated [35].

Suprainfection has strong implications in evolutionary biology. First, it supports the study of complementation in experimental evolution [82]. Second, it raises the question of the origin of the dependence: is the dependent genotype the descendent of a defective mutant of the self-sufficient genotype or is it a fully functional mutant that has become parasitic of another genotype? Finally, one can consider the evolution of suprainfection towards ambinfection if, for instance, the self-sufficient genotype undergoes deleterious mutations that are complemented by the other genotype.

**Ambinfection** Two parasite genotypes define an ambinfection when they can only infect together. This can result from an association between a genotype that does not grow when alone and another genotype that provokes explosive infections when alone: once together, the latter facilitates the reproduction of the former, while the former inhibits the growth of the latter. It can also derive from the association between two genotypes that provoke explosive infections when alone but inhibit each other when together so that their parasite loads become bounded. Finally, it can result from the crossfacilitation of two genotypes that cannot grow alone otherwise, that is obligate mutualism (results not shown).

This last situation often occurs in multipartite plant viruses. These viruses, such as the bipartite *Tomato golden mosaic virus* (TGMV) [83], have a fragmented genome; they encapsidate each fragment in a separate capsid and, typically, all genomic fragments need to be inoculated in the same host to produce a viable infection.

Plasmids causing bacteria to be virulent offer another example of ambinfection. For instance, the bacterial agent of bubonic plague, *Yersinia pestis*, relies on a set of 12 proteins encoded by a 70-kb plasmid (called pYV) to inhibit phagocytosis and hence proliferate inside lymph nodes in humans [84]. The competitive Lotka-Volterra framework is however less relevant in these cases because it relies on the physical independence between competing individuals and may not suit to intracellular molecules such as plasmids (nor hyperparasitism).

The fact that parasites causing ambinfections cannot be detected separately may lead researchers to consider such organisms as a single infectious agent and perhaps explains why this pattern is rarely found.

### Perspectives

We argue that it is necessary to challenge the coinfection/superinfection dichotomy for several reasons. First, the inoculation outcome can also depend on the initial conditions and not only on the parasite traits (the difference between superinfection and priorinfection). Second, parasite load explosion in finite time does not fall in either of these two patterns nor ‘in between’. Third, even if priorinfection and ultrainfection are added to the dichotomy, there remains an ambiguity between ‘coinfection’ with the two single infections (what we call coinfection) and ‘coinfection’ with one or no single infection (ambinfection and suprainfections) for instance. The typology we introduce is less ambiguous and manages to capture infection patterns of actual biological systems (see examples above) that are otherwise unclassifiable.

Our results stress that if only prevalence data are available, it is impossible to distinguish between infection patterns because this requires knowledge about the within-host dynamics (or at least the outcome of cross-inoculations). As shown in appendix Table S1-10 and formalised in appendix E.2, priorinfection, ultrainfection and the two superinfections are epidemiologically equivalent as all the classes can be observed except for the doubly infected one. Leaving aside the epidemiology and observed prevalences, these patterns are shaped by different underlying mechanisms and may therefore have distinct evolutionary pathways.

Synergy between field or experimental work and theoretical analysis emerges as a necessity for studying multiple infections. Models inspire new hypotheses to test on natural systems, such as shifts of infection patterns (see e.g. [65]), and direct observations help to improve frameworks, as in the case of phage mutual exclusion [24], which does not fit in the super/co-infection dichotomy. In this sense, our within-host model could be tailored to a particular biological system by integrating specific mechanisms and explicit host immune response in order to better delineate the infection pattern conditions. A particular improvement would be to introduce frequency-dependence, which notably arises in non-enzymatic and consumable public productions, and therefore look for the new infection patterns the model generates but also the lost ones. Nonetheless we believe that the present typology of infection patterns for two genotypes is sufficient to describe most natural situations.

The outcome of a particular inoculation experiment could also be altered by environmental effects, host heterogeneity or stochasticity. Environmental effects and host heterogeneity can be addressed by splitting the host population into as many groups as needed for having constant environmental conditions and host homogeneity within these groups. Then the infection pattern has to be independently identified in each group to determine whether it is environment or host dependent. For example the above procedure could be used to determine whether the outcome of a given set of inoculation experiments depends on an environmental factor, such as temperature, or on host genotype. Stochasticity will not affect the nature of infection patterns, only the probability under which each may be observed. The frequency of each outcome could thus be used to draw a probability distribution of the infection patterns.

Our results may help to identify the infection pattern of a natural system but also to better model the associated epidemiological dynamics by choosing the right flows between compartments and write the appropriate equations, thus providing further insight into the underlying within-host interactions. We acknowledge that this approach is still limited because it assumes there are at most two genotypes, while genotypic diversity is often higher [5]. Fully determined models of within-host growth with multiple interactions are rare. First, because of obvious practical constraints [55] but also because it is already possible to draw key qualitative conclusions, as attested by the popularity of the co-infection models. Furthermore, most evolutionary frameworks such as adaptive dynamics consider no more than two different genotypes, the resident and one mutant [85].

Multiple infections are highly polymorphic and polyspecific in nature, where we expect to find many more infection patterns than those listed here. A third genotype could for instance allow two genotypes that are otherwise in a priorinfection setting to coexist by inhibiting both of them. Our typology is helpful to start investigating higher rank infection patterns. Indeed, if we consider three genotypes, there are exactly four possible partitions of the set of genotypes ({1,2,3} & {∅},{1} & {2,3},{1, 2} & {3} and {1, 3} & {2}). Within each of these partitions, non empty sets of genotypes can be seen as “supergenotypes”. The superinfection pattern associated to the supergenotype - genotype couple (or supergenotype alone in the case of the first partition) can then be determined as in our model, with the additional convention that for a supergenotype to be considered as having reached stationary infection, all genotypes that belong to it must have done so. The whole infection pattern is finally provided by the complementary analysis of these four supergenotype - genotype infection patterns plus the three infection patterns of the genotype-genotype couples. Therefore, higher rank infection patterns can be iteratively described by the infection patterns given for two genotypes only.

We have drawn a map of multiple infections along with a clear formalism, thus filling a gap left by previous frameworks and shedding new light on patterns that are observed empirically. Our infection patterns framework can easily be used to study the genetic diversity of microbiota and other commensal and/or mutualistic communities. Furthermore, if we consider non parasitic species and patches of habitat instead of hosts, our results can be extended to the field of meta-community ecology.

## Acknowledgements

The authors thank Gael Thebaud, Remy Froissart, Stephane Blanc for their helpful suggestions of biological examples of infection patterns and Yves Dumont and Matthieu Hillairet for their useful expertise in system dynamics.

## Authors’ Contributions

Conceived the project: M. T. Sofonea, S. Alizon and Y. Michalakis.

Performed the analysis: M. T. Sofonea.

Wrote the article: M. T. Sofonea, S. Alizon and Y. Michalakis.

